# Very long-chain ceramides in muscle associate with insulin resistance independent of obesity

**DOI:** 10.1101/2025.03.04.641442

**Authors:** Søren Madsen, Harry B Cutler, Kristen C Cooke, Meg Potter, Jasmine Khor, Christoph D Rau, Stewart WC Masson, Anna Howell, Zora Modrusan, Anthony S Don, Jacqueline Stöckli, Alexis Diaz Vegas, David E James

## Abstract

Lipids, in particular ceramides and diacylglycerols (DAGs), are implicated in insulin resistance, however their precise roles remain unclear. We leverage natural genetic variation to examine muscle lipids and systemic insulin resistance (IR) in 399 Diversity Outbred Australia mice. Adipose mass was associated with 55% of muscle lipids and IR, with DAGs as the only enriched lipid class. To disentangle adiposity and muscle lipid contributions, we used two approaches: (1) linear modelling of muscle lipids corrected for adipose mass on systemic IR, and (2) stratifying mice into insulin sensitivity quartiles within adiposity bins. Both revealed that very long-chain ceramides, but not DAGs, were linked to IR. Transcriptomic and proteomics further associated these ceramides with cellular and mitochondrial stress. DAGs correlated with leptin expression in muscle, suggesting they originate from muscle-residing adipocytes. We propose that many muscle lipids, including DAGs, associate with IR due to adipose accumulation rather than directly influencing muscle insulin sensitivity. By addressing the relationship between adiposity and metabolic state, we identified very long-chain muscle ceramides as highly associated with IR independently of adiposity.

## Introduction

Insulin resistance (IR), defined by a defect in insulin-regulated glucose metabolism, is one of the earliest risk factors for chronic cardiometabolic disorders and is shaped by both genetic and environmental factors (James *et al*, 2021). At a systemic level, IR triggers a compensatory increase in insulin secretion from pancreatic β-cells, which helps maintain stable blood glucose levels despite the presence of IR (James *et al*, 2021). This compensatory hyperinsulinemia has been associated with several pathologies, including cancer, heart disease, and metabolic associated fatty liver disease (Zhang *et al*, 2023; Steneberg *et al*, 2015; Després *et al*, 1996).

Skeletal muscle plays a crucial role in systemic insulin sensitivity, as it is the primary site for postprandial carbohydrate uptake (Thiebaud *et al*, 1982). Moreover, increased muscle insulin sensitivity is sufficient to protect mice against diet-induced metabolic dysfunction, such as fasting hyperinsulinemia and hyperglycemia (Tsao *et al*, 1996; Beckerman *et al*, 2021; Diaz-Vegas *et al*, 2024a; Nelson *et al*, 2022). At the molecular level, muscle IR has been associated with excessive lipid accumulation within the tissue, a process known as lipotoxicity. This results in an increase in specific lipid intermediates, such as diacylglycerols (DAGs) and the sphingolipid ceramide, which are proposed to decrease insulin-induced GLUT4 translocation to the cell surface (Perreault *et al*, 2018; Erion & Shulman, 2010; Turpin-Nolan *et al*, 2019). However, the relative importance of these lipids for IR is still debated (Erion & Shulman, 2010; Chaurasia & Summers, 2021; Zierath, 2007).

The cause of DAG accumulation in skeletal muscle IR remains unclear, but it is thought to result from an imbalance between fatty acid uptake and oxidation (Koves *et al*, 2008). Excess fatty acid supply overwhelms mitochondrial oxidative capacity, diverting fatty acids towards *de novo* DAG synthesis, leading to cellular accumulation and contributing to IR (Morino *et al*, 2006; Kraegen *et al*, 2009). Interestingly, *in vivo* studies using labelled tracers suggest that *de novo* DAG synthesis alone does not fully account for this accumulation (Perreault *et al*, 2018), pointing to other pathways, such as phospholipid degradation, changes in the interconversion between sphingomyelin/ceramide, and direct delivery of DAGs by plasma lipoproteins (Perreault *et al*, 2018). Plasma lipoproteins, especially chylomicrons and very low-density lipoproteins (VLDL), can transport various lipid species, including DAGs, in their core or on their surface (Cohen & Fisher, 2013). These lipoproteins deliver a significant portion of their lipid content to muscle tissue, potentially contributing to a substantial amount of DAG accumulation, particularly in the context of obesity and IR (Kim *et al*, 2001). Furthermore, adipocyte infiltration in skeletal muscle has been observed across various models of obesity (Jani *et al*, 2021; Hilton *et al*, 2008; Wang *et al*, 2024), potentially altering muscle lipid composition and contributing to IR.

Ceramide accumulation in skeletal muscle has also been strongly linked to IR (Chaurasia & Summers, 2021). The primary mechanism driving ceramide accumulation in muscle is the *de novo* synthesis pathway, which begins with the condensation of serine and palmitoyl-CoA by the enzyme serine palmitoyltransferase - the rate-limiting step in this process (Larsen & Tennagels, 2014). Additional pathways increasing muscle ceramides include sphingomyelin hydrolysis and the salvage pathway, which are cell-autonomous processes (Chaurasia & Summers, 2021). These pathways are also linked to muscle IR. For example, the salvage pathway for ceramide production is activated by chronic inflammation, a condition commonly seen in individuals with obesity (Green *et al*, 2021). Furthermore, selective activation of sphingomyelin hydrolysis is sufficient to drive IR in rat myotubes (Diaz-Vegas *et al*, 2023).

Recent advancements in mass spectrometry-based global lipidomics have enhanced our understanding of muscle lipotoxicity. These analyses have highlighted the complexity of lipid species, their subcellular distribution and established correlative links between muscle lipid composition and the development of IR in mice and humans (Turpin-Nolan *et al*, 2019; Perreault *et al*, 2018). While these studies highlight links between lipid species and IR, fully elucidating this relationship in the context of obesity remains challenging, particularly due to the genetic variance influencing IR development (James *et al*, 2021; Hunter, 2005; Nelson *et al*, 2022). Recent studies using genetically diverse mouse populations have significantly advanced our understanding of metabolic diseases (Masson *et al*, 2023; Chella Krishnan *et al*, 2023; Churchill *et al*, 2012; Nelson *et al*, 2022; van Gerwen *et al*, 2024). These models provide controlled environments, access to various biological tissues, and a broad range of phenotypic diversity, making them highly valuable for investigating complex diseases (Masson *et al*, 2024). A prominent example is the Diversity Outbred Australia (DOz) mouse population, which exhibits remarkable phenotypic variation even in younger animals fed a healthy chow diet (Masson *et al*, 2023).

Here, we utilised the DOz mouse population to investigate the link between lipotoxicity in skeletal muscle and systemic IR across natural phenotypic variation by combining metabolic phenotyping with paired lipidomics, proteomics and transcriptomics. While DAGs correlated most strongly with IR, this relationship was dependent on adipose tissue mass, suggesting they may not directly regulate muscle insulin sensitivity. In support of this, DAGs were closely linked to leptin gene expression in skeletal muscle, an adipocyte specific transcript, indicating adiposity and possibly adipocyte infiltration as a major contributor to DAG accumulation in skeletal muscle. In contrast, very long-chain ceramides correlated with systemic IR independently of obesity and were strongly associated with molecular markers of mitochondrial stress, highlighting their potential role in skeletal muscle metabolic dysfunction.

## Methods

### Animal experiments

Experiments were performed in accordance with NHMRC guidelines and under approval of the University of Sydney Animal Ethics Committee, approval numbers #1274 and #1988. All mice were maintained at 23°C on a 12 hr light/dark cycle.

### Animal experiments

Male DOz mice were bred and housed at the Charles Perkins Centre, University of Sydney, NSW, Australia, as detailed in (Masson *et al*, 2023). Animals were 10-12 wks of age at the start of experiments and had free access to a standard laboratory chow diet containing 16% calories from fat, 61% calories from carbohydrates, and 23% calories from protein (Specialty Feeds) or an in-house made high-fat diet (HFD) containing 60% calories from fat, 20% calories from carbohydrate, and 20% calories from protein (25% g casein, 0.4% g L-cystein, 16% g cornstarch, 9.3% g sucrose, 6.4% g cellulose, 31.3% g lard, 3.2% g safflower oil, 6.4% g AIN-93 mineral mix [MP Biomedicals], 1.3% g AIN-93 vitamin mix [MP Biomedicals], 0.3% g choline. Total adipose tissue and lean mass were measured with EchoMRI-900 (EchoMRI Corporation Pte Ltd, Singapore) at 14 wks of age (for chow) and at 8 or 24 wks (for HFD diet)). Glucose tolerance was determined by a glucose tolerance test at the same age by fasting mice for 6 h (from 07am) before oral gavage of 20% glucose solution in water at 2 mg/kg lean mass. Blood glucose concentration was measured by glucometer (Accu-Chek, Roche Diabetes Care, NSW, Australia) from tail blood 0, 15, 30, 45, 60, and 90 min after oral gavage of glucose. Blood insulin levels at the 0 and 15 min timepoints were measured by mouse insulin ELISA Crystal Chem USA (Elk Grove Village, IL) according to the manufacturer’s instructions. Homeostatic Model Assessment for IR (HOMA-IR) was calculated as follows:

‘Insulin (ng/mL)’ / (5803*10^6^)*10^12^/6 x ‘Blood glucose (mmol/L)’*18 / 405

Mice were sedated by pentobarbital (⅕ dilution) and quadriceps muscle removed and snap frozen in liquid nitrogen following cervical dislocation.

### Quadriceps muscle lipidomics Lipid extraction

Lipid extraction was performed as previously reported (Turner *et al*, 2018). Briefly, a two-phase lipid extraction from frozen quadriceps muscle (∼20 mg) was performed using the methyl-tert-butyl ether (MTBE)/methanol/water method. Frozen tissue was powdered in liquid nitrogen, transferred to 2 mL Eppendorf tubes, and homogenised in 0.2 mL methanol containing 0.01% butylated hydroxytoluene (BHT) using a sonicator water bath (Thermoline, model 505) in a cold room. The homogenates were spiked with an internal standard mixture consisting of 5 nmoles 19:0/19:0 phosphatidylcholine (PC) and d7-cholesterol, 2 nmoles 18:1/15:0 d7-diacylglycerol, d18:1/17:0 glucosylceramide, 17:17:0 phosphatidylethanolamine (PE), and d18:1/17:0 SM, 4 nmoles 14:0/14:0/14:0/14:0 cardiolipin, 500 pmoles d18:1/17:0 ceramide, and 200 pmoles d17:1 sphingosine and d17:1 S1P. Next, 1.7 mL MTBE was added, and the samples were sonicated for 30 minutes in an ice-cold water bath (Thermoline Scientific, Australia). Phase separation was induced by adding 417 µL of mass spectrometry-grade water, followed by vortexing at maximum speed for 30 seconds and centrifugation at 1000 x g for 10 minutes. The upper organic phase (∼600 µL) was transferred into a 96-well deep plate. The aqueous phase was re-extracted three times using the same MTBE/methanol/water mixture, and the organic phases were combined in the deep-well plate (∼1800 µL). The organic phase was then dried under vacuum using a Savant SC210 SpeedVac (Thermo Scientific). The dried lipids were resuspended in 500 µL of 80% methanol/0.2% formic acid/2mM ammonium formate, mixed at 2000 rpm for 5 minutes, and stored at −20°C until further analysis.

### Lipid detection

Lipids were quantified using selected reaction monitoring (SRM) on a TSQ Altis triple quadrupole mass spectrometer, coupled with a Vanquish HPLC system (ThermoFisher Scientific). Lipid separation was achieved on a 2.1 x 100 mm Waters Acquity UPLC C18 column (1.7 µm pore size) with a flow rate of 0.28 mL/min. The mobile phase consisted of two components: phase A (0.1% formic acid and 10 mM ammonium formate in 60% acetonitrile/40% water) and phase B (0.1% formic acid and 10 mM ammonium formate in 90% isopropanol/10% acetonitrile). The total run time was 25 minutes, starting with 20% phase B for the first 3 minutes, then gradually increasing to 100% phase B from 3 to 14 min. The system was held at 100% phase B from 14 to 20 min, returned to 20% phase B by 20.5 min, and held there for an additional 4.5 min. Ceramides, sphingomyelin, sphingosine, and sphingosine-1-phosphate were identified using the [M+H]+ precursor ion, with product ions at *m/z* 262.3 (sphinganine), 264.3 (sphingosine), 266.3 (sphinganine), or 184.1 (sphingomyelin). Diacylglycerols were identified by the [M+NH4]+ precursor ion, with a product ion corresponding to the neutral loss of a fatty acid +NH3. Cardiolipins were identified by the [M+H]+ precursor ion and the product ion corresponding to the neutral loss of a diacylglycerol. Cholesterol was detected using a precursor ion at *m/z* 369.4 and a product ion at *m/z* 161.1. We measured glycerolipids (diacylglycerols; DAGs), glycerophospholipids (phosphatidylethanolamines; PEs, phosphatidylcholines; PCs, cardiolipin; CLs, and plasmalogen phosphatidylethanolamines; PEPs), cholesterol and sphingolipids (sphingosines; Sph, sphingosine-1-phosphates; S1Ps, sphingomyelins; SMs, Ceramides; Cers, and Hexosylceramides; Hexs) in quadriceps from all mice (Supplemental table 1).

### Lipid quantification

Lipids were quantified by calculating the area under the peak for a specific precursor–product ion pair. Each peak contained at least 8–10 scans for that ion pair. Peak detection and integration was performed using TraceFinder. To determine the amount of each lipid, the ratio to its class-specific internal standard was multiplied by the amount of internal standard added, then divided by the mass of extracted tissue. A detailed description of these analyses can be found in (Diaz-Vegas *et al*, 2024b).

### Transcriptomics

We randomly selected 92 chow fed animals for transcriptomic analysis. RNA from mouse quadriceps was extracted using a combined TRIzol/chloroform/RNeasy kit (Invitrogen/Sigma Aldrich/Qiagen) spin column protocol. Muscles were pulverised in liquid nitrogen and lysed in 0.5 mL Trizol with a Mixer Mill (MM400, Retsch) at room temperature and incubated for 5 min. 0.1 mL chloroform was added, tubes inverted ∼10 times and left at room temperature for 3 min, then spun at 14.000 g at 4C for 10 min. The supernatant was transferred to new tubes containing 0.3 mL 70% ethanol. The sample:ethanol mix was added to the RNeasy spin columns and RNA was extracted following the manufacturer’s instructions. RNA integrity was assessed by 4200 TapeStation System (Agilent). All samples had RNA Integrity Number equivalent greater than 8. KAPA mRNA HyperPrep Kit (Roche) was used for RNA sequencing library preparation following manufacturer’s instructions with 500 ng of RNA as input and sequenced on NovaSeq 6000 (Illumina). Quality of sequencing data was assessed with multiQC (Ewels *et al*, 2016). Gene expression levels were quantified with Kallisto (Bray *et al*, 2016) version 0.50.1 with the following flags: kallisto quant-b 100--single-l 50-s 20. Mouse index file was downloaded from (November 2023): https://github.com/pachterlab/kallisto-transcriptome-indices/releases. Transcript levels were estimated with the r package tximport (Soneson *et al*, 2015) and EdgeR (Robinson *et al*, 2010) was used to filter lowly expressed genes. No samples were excluded. Raw sequencing data have been deposited at Gene Expression Omnibus (accession number GSE290031 - reviewer token: qfybwmkwnlwjzsb).

### Proteomics

Proteomics data has been published previously (Masson *et al*, 2023) and has been deposited on ProteomeXchange (PXD042277). The data consisted of two fractions for each quadriceps muscle; a mitochondria enriched fraction and the non-mitochondrial fraction from 228 chow fed animals. The overlap with the lipidomics data was 197 animals. Proteomics raw data files were researched using DIA-NN (v1.8) using a library-free FASTA search against the reviewed UniProt mouse proteome (downloaded March 2023). The protease was set to Trypsin/P with one missed cleavage, N-term M excision, carbamidomethylation, and M oxidation options on. Peptide length was set to 7–30, precursor range 350–1650, and fragment range 300– 2000, and false discovery rate (FDR) set to 1%. Both the mitochondrial enriched and non-mitochondrial samples were filtered based on the mitocarta 3.0 and that a protein was quantified in at least 50% of samples. Protein intensities were log2 transformed and median normalized to account for technical variation in total protein levels across all samples.

### Data analysis

All data analysis and visualisation were performed in R (version 4.1 - 4.3). Pearson correlation was calculated with cor.test function in base R and bicor calculated with WGCNA package (Langfelder & Horvath, 2008). To determine the association between a lipid feature and HOMA-IR, we used the following linear model:

Lipid feature ∼ α + β1 HOMA IR x β2 diet + β3 TimeOnDiet + ε fitted with the glm function. The type II sum of squares was calculated using the ANOVA function of the R car package. To correct lipid features for adiposity, we built a linear model for lipid feature ∼ adipose tissue mass + ε with lm base R function and worked with the residuals (ε).

In general, all reported p-values were corrected for multiple testing using the false-discovery rate correction (p.adjust(p-value, method = “fdr”)) with one exception (the HOMA-IR quartile analysis).

For the segmentation of animals in HOMA-IR quartiles across adiposity bins, animals were divided into 10-percentage-points bins starting from 15% adiposity (± 5 percentage points) and ending at 40%. This cut-off was chosen to ensure we had sufficient mice for the quartile analysis. We compared lipid features between lowest (Q1) and highest (Q4) HOMA-IR quartiles across the whole adiposity span with increments of 1 percentage points by t-test on log(x+1) transformed lipidomics data following q-value estimation by the fdrtool function in fdrtool package (Strimmer, 2008). To test for mean difference of lipid features between Q1 and Q4 we applied one-sample t-tests to all log2 fold-changes across all adiposity bins with the null hypothesis: log2 fold change = 0 to assess if the average lipid abundance in Q4 is different to Q1. P-values were adjusted with the Bonferroni method.

Gene set enrichment analysis was performed with ClusterProfiler (Wu *et al*, 2021) and ReactomePA (Yu & He, 2016).

## Results

### Outbred mice show significant phenotypic diversity

399 male DOz mice were fed high-fat diet (HFD) or standard chow for varying durations (Fig. 1A): 1) standard chow (n = 203); 2) HFD for 8 wks (n = 80) or for 15 wks (n = 116). Overall, we observed significant phenotypic diversity in all metabolic traits (Fig. 1B-E). Adipose tissue mass (Fig. 1B) and fasting insulin (Fig. 1C) both varied ∼250-fold from lowest to highest, and the Homeostatic Model Assessment for IR (HOMA-IR), a readout of systemic insulin sensitivity, varied 450-fold (Fig. 1D). Both diet and duration of feeding impacted all metabolic traits with the longer term HFD group having a significantly higher increase in adipose tissue mass, fasting insulin and HOMA-IR compared with the short term HFD group, which again was increased compared to the chow group (Fig 1B-D). Fasting blood glucose was increased with HFD, but no difference was seen between the two HFD groups (Fig. 1E). Fasting insulin was the main contributor to HOMA-IR, explaining 92% of the variance in HOMA-IR (Fig. 1F). Overall, DOz mice capture a huge phenotypic range of metabolic parameters.

**Figure 1.**
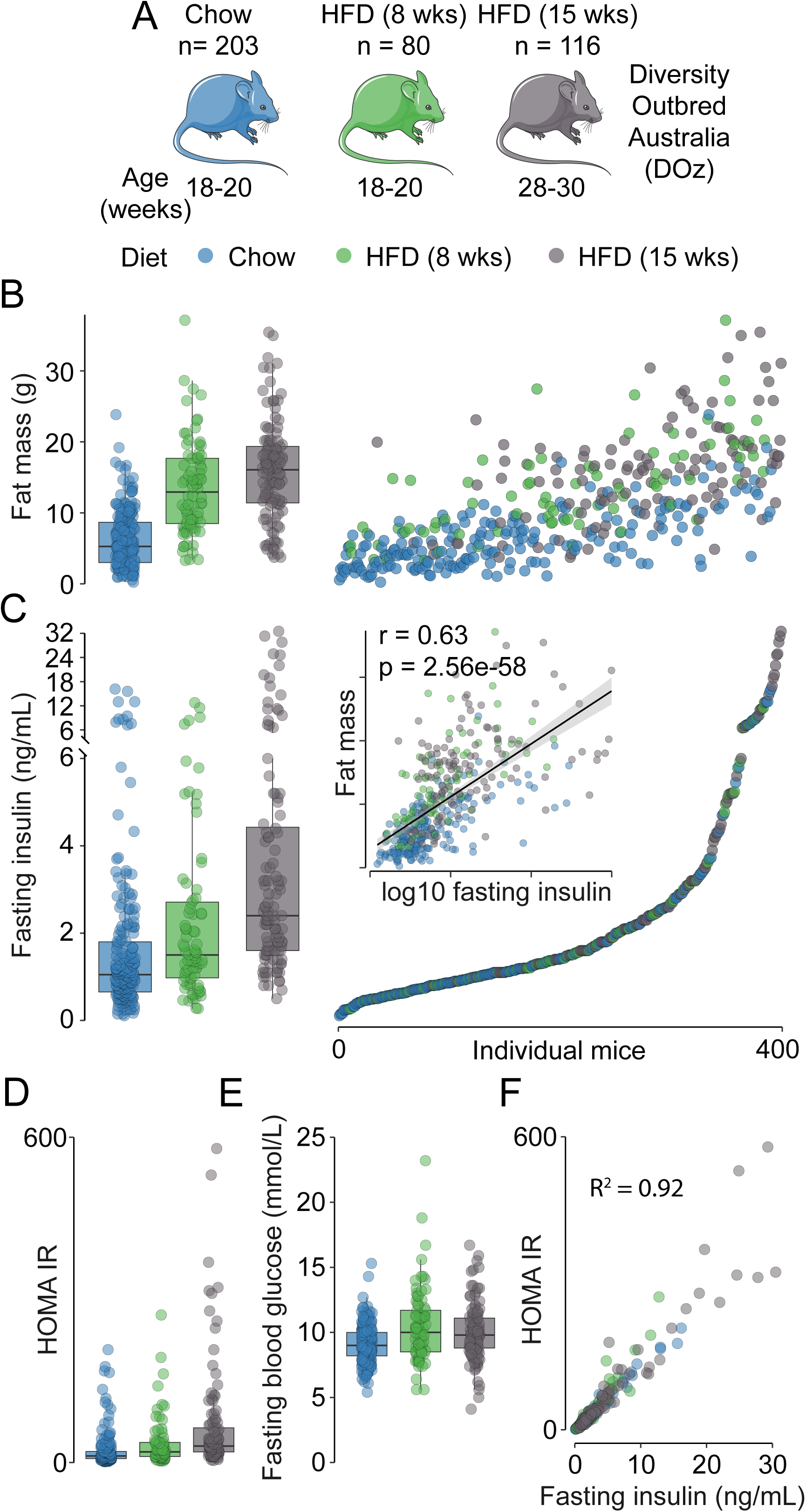
Metabolic traits across 399 Diversity Outbred Australia (DOz) mice fed high-fat diet or chow. **A)** Male DOz mice were either fed chow (n = 203) or a high-fat diet for 8 (n = 80) or 15 weeks (n = 116). The chow fed group was aged-matched to the 8 weeks HFD group. **B & C)** Fat mass **(B)** and fasting blood insulin **(C)** across all mice. Boxplots show traits grouped by diet and scatter plots show each trait ordered by blood insulin (lowest to highest). Insert shows the relationship between fat mass and blood insulin. For both traits, chow < 8 weeks < 15 weeks significantly different. **D)** HOMA-IR across diet groups; chow < 8 weeks < 15 weeks significantly different. **E)** Fasting blood glucose across diet groups; chow group significantly different from HFD groups only. **F)** Relationship between fasting blood insulin and HOMA-IR.

### Targeted lipidomics reveals that muscle diacylglycerols are associated with systemic insulin sensitivity

Because of the strong link between lipotoxicity and IR we employed a targeted lipidomic strategy to identify specific lipid species in quadriceps muscle that were associated with insulin sensitivity. Of the lipids measured by our targeted approach, phosphatidylcholines (PCs) (38%), cholesterol (29%) and phosphatidylethanolamines (PEs) (21%) made up 87% of the total lipids measured (Fig. S1A).

As with the metabolic parameters, muscle lipids varied greatly across the whole cohort (Fig. 2A). Dimension reduction revealed good clustering of the lipidome with chow and the longer term HFD group forming two distinct clusters and the short term HFD group was intermediate (Fig. 2B). Correlation analysis demonstrated that adipose tissue mass had a strong influence on the muscle lipid profile, as 55% of identified lipids had a significant association with this parameter (Fig. 2C). Overall, muscle DAGs showed a strong tendency toward overrepresentation in associating with adipose tissue mass (9 out of 11 DAGs quantified, p = 0.081, Fisher’s exact test) indicating that DAG levels in skeletal muscle may simply depend on the state of obesity. Additionally, muscle DAGs were the only lipid class that were overrepresented in their association to fasting insulin (p = 5e-4), fasting blood glucose (p = 8e-4) and systemic insulin sensitivity (HOMA-IR, p = 9e-4) (Fig. 2C) with sphingomyelins (SMs) overrepresented in the association with fasting insulin only (p = 0.02). DAGs containing unsaturated fatty acids with few double bonds (18:1-18:1, 16:0-18:1 and 18:1-18:2) were positively associated with these traits, whereas highly polyunsaturated DAGs (16:0-22:6 and 18:0-22:6) were inversely associated. Of note, ceramides showed an inverse relationship to adipose tissue mass, as nine of the 14 ceramide species measured were negatively associated with adipose tissue mass. The only ceramide to associate with fasting insulin and HOMA-IR was the C18 ceramide d18:2-18.0 although the association was weak (r = 0.16, p = 7e-3).

**Figure 2.**
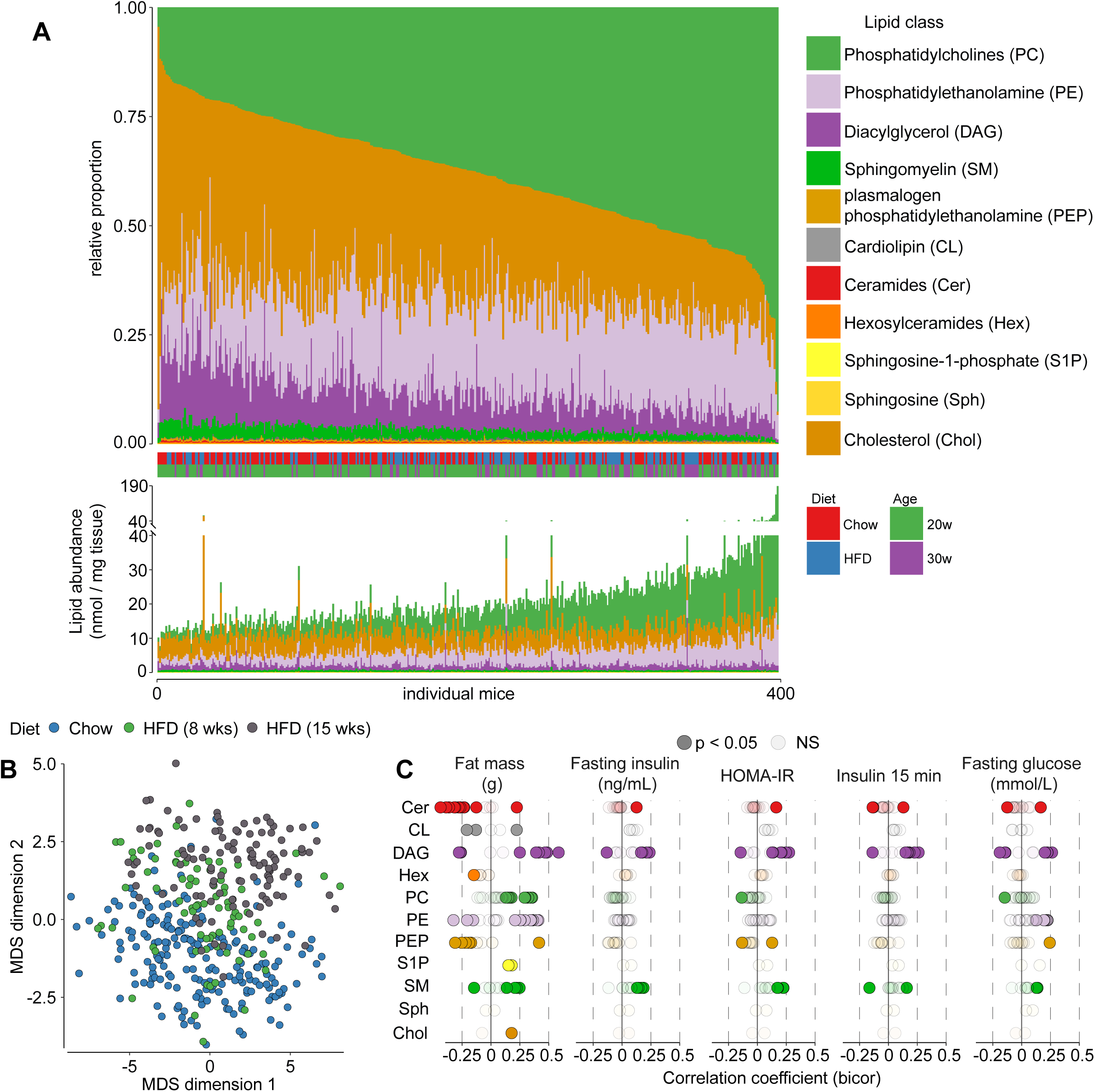
Target lipidomics in quadriceps from 399 DOz male mice. **A)** Relative (top) and absolute (bottom) abundance of lipid classes quantified by targeted lipidomics. Both are ordered by abundance of absolute total phosphatidylcholine abundance (lowest to highest). **B)** Dimension reduction by multidimensional scaling (MDS) of all lipid features. **C)** Bicor correlation coefficient of all lipid features to fat mass, fasting blood insulin, HOMA-IR, blood insulin at 15 min during glucose tolerance test and fasting blood glucose.

Overall, these data show that there is considerable variance in the skeletal muscle lipidome between individual DOz mice fed either chow or HFD, and the predominant source of variation in lipid levels appears to stem from differences in adiposity.

### Ceramides are associated with systemic insulin resistance independently of adipose tissue mass

Given that adiposity is strongly associated with IR and the strong link between adiposity and DAGs in muscle, this prompted us to investigate whether adipose tissue mass mediated the observed relationship between DAGs and systemic insulin sensitivity. To do this, we used linear modelling to examine associations between systemic insulin sensitivity, measured by HOMA-IR, and muscle lipids independently of other variables such as diet or adipose tissue mass. Consistent with the previous correlation analysis, the lipids most strongly associated with HOMA-IR after adjusting for diet were DAGs (18:1-18:1, 16:0-18:1, and 18:1-18:2), the C18 ceramide (d18:2-18:0), and a polyunsaturated DAG (18:1-20:4) (Fig. 3A). Furthermore, two SMs were significantly associated with HOMA-IR.

**Figure 3.**
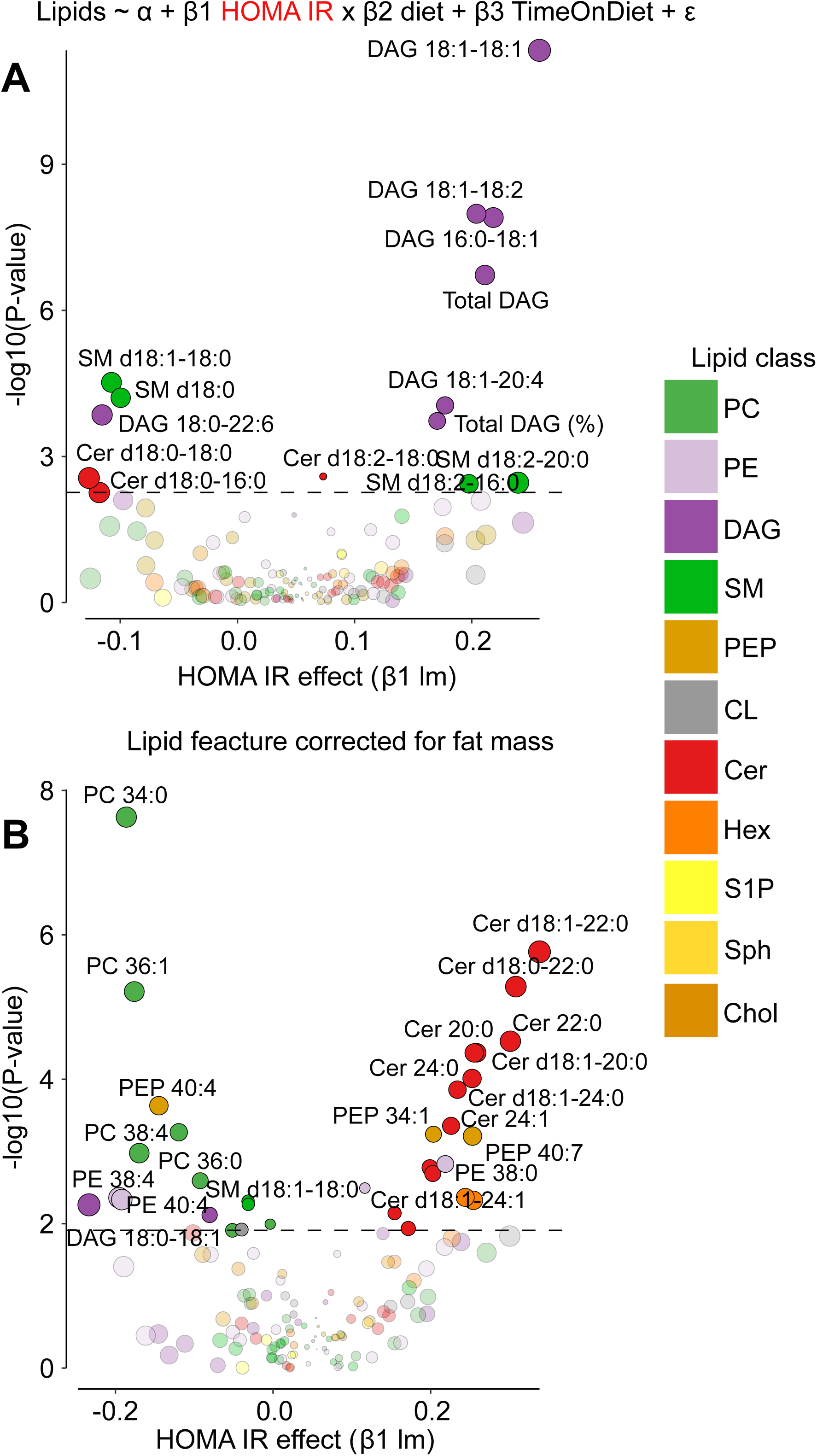
Linear modelling showing relationship between HOMA-IR and each lipid feature. **A)** Main effect of HOMA-IR on individual lipid features from linear models accounting for diet and time of diet. **B)** Main effect of HOMA-IR on individual lipid features corrected for overall fat mass from linear models accounting for diet and time of diet. Size of data points are sized to the absolute change in HOMA-IR.

Interestingly, lipids that were associated with increased whole-body insulin sensitivity (decreased HOMA-IR) included long-chain SMs such as d18:1-18:0, as well as DAG species like 18:0-22:6. Intriguingly, long-chain ceramides such as d18:0-16:0 and d18:0-18:0 were also positively linked to insulin sensitivity (Fig 3). Since our primary focus was on identifying muscle lipids with associations to IR, and since insulin sensitivity and muscle lipids were influenced by adiposity, we constructed a model where each lipid feature was corrected for overall adipose tissue mass. This allowed us to analyse the muscle lipidome and its association with IR independently of adipose tissue mass. Strikingly, this revealed that very long-chain ceramides (d18:1-22:0, d18:1-20:0, d18:1-24:0, d18:0-22:0, d18:0-20:0, d18:1-24:1) were all associated with increased HOMA-IR (Fig. 3B). In addition to these ceramides, the very long-chain hexosylceramide (Hex) species, d18:1-24:1 and d18:1-24:0, both derived from very long-chain ceramides, were also associated with HOMA-IR. Finally, both PCs and PEs, which are the primary phospholipids in cellular membranes, were positively associated with insulin sensitivity.

### Stratified quartile-based assessment of muscle lipids by HOMA-IR across the full range of adiposity

To further explore how muscle lipids relate to whole-body insulin sensitivity independently of adiposity, we stratified mice into adiposity bins of 10-percentage-point increments and categorized them into HOMA-IR quartiles within each bin (Fig. 4A & B). HOMA-IR was significantly different in the most insulin sensitive quartile (Q1) and the most IR quartile (Q4) across all bins, creating groups with similar adiposity but distinct insulin sensitivity. This analysis emphasises the strength of using DOz mice as in individual inbred mouse strains like C57BL/6J mice whole-body IR is always associated with increased adiposity (Nelson *et al*, 2022). Bin sizes ranged from 30 to 58 animals (Fig. 4B). We examined the lipids between Q1 and Q4 in each adiposity bin with a particular focus on those lipids that were most strongly associated with IR (Fig. 3). Two ceramides were nominally elevated in all but one bin and significantly increased in 50% of the bins, while DAGs showed no consistent changes across the bins (Fig. 4C). This is consistent with the hypothesis that DAGs are associated with systemic IR as a consequence of increased adiposity, whereas ceramides seem more intrinsic to the muscle itself.

**Figure 4.**
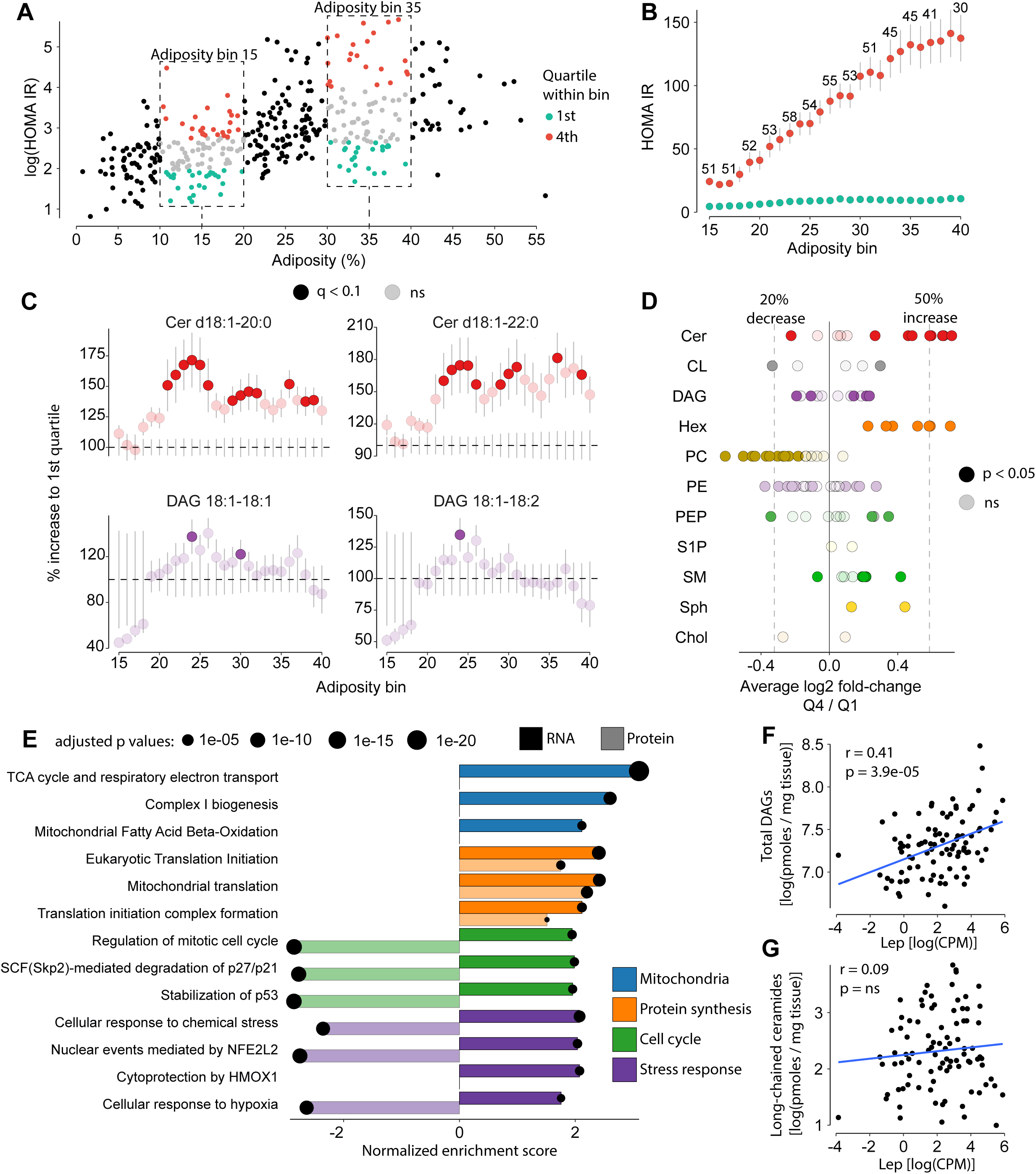
Stratified quartile-based assessment of muscle lipids by HOMA-IR across adiposity bins. **A)** Relationship between adiposity and HOMA-IR. Two adiposity bin of 10 percentage points highlighted. Within each bin, 1st and 4th quartile (Q) of HOMA-IR are shown. **B)** HOMA-IR across all adiposity bins. Number above indicates the total number of animals (Q1 + Q4). Significant difference between Q1 and Q4 in all adiposity bins. **C)** Comparison of ceramides (Cers) and diacylglycerols (DAGs) abundance between Q1 and Q4 relative to Q1 across all adiposity bins. **D)** Overall average difference between Q1 and Q4 across all adiposity bins. Significance tested by one-sample t-test (null hypothesis: log2 fold-changes centers around 0 (log2FC = 0)). **E)** Selected pathways associated with very long-chain ceramides from both transcriptomic (n = 92) and proteomic (n = 197) data. **F)** The relationship between total DAG abundance and leptin gene expression in skeletal muscle. **G)** The relationship between very long-chain ceramides and leptin gene expression. Error bars are standard error over mean (SEM).

To expand this type of analysis to all lipid features, we performed one-sample t-tests on the log2 fold-changes across all bins to determine if they differed from 0. Overall, this showed that 10/14 ceramides were significantly different between Q1 and Q4 groups (9 increased with systemic IR), whereas only 5/11 DAGs (3 increased with IR) were different (Fig. 4D). Consistent with previous analysis (Fig. 3), the very long-chain ceramides (20, 22 and 24 carbons) showed the largest increase in the insulin resistant group (>50% increase). Furthermore, all Hex and Sph species quantified in our data and 7/11 SMs were also significantly increased. We do note, as mentioned above, that three DAGs were significantly increased in the insulin resistant groups across all adiposity bins, however the average increase was <20% and the profile of change across the adiposity bins was less consistent (Fig. S2A). Furthermore, C18 ceramides, the muscle ceramide class often associated with impaired metabolic state (Hammerschmidt & Brüning, 2022), were unchanged across all bins. Together, this shows that very long-chain ceramides and sphingolipids in general were the lipids that were most significantly and consistently increased in quadriceps muscle from insulin resistant animals and this occurred independently of adiposity.

### Molecular signatures of selected muscle lipid features

Ceramides have been strongly associated with several cellular pathways including inflammation, oxidative stress and mitochondrial dysfunction (Chaurasia & Summers, 2021). To understand the molecular underpinnings of elevated ceramide levels in skeletal muscle, we quantified the quadriceps transcriptome (Supplemental table 2) and proteome (Supplemental table 3) from a subset of chow fed mice and correlated these to the total abundance of all very long-chain ceramides elevated in the IR state. From the transcriptome, we found four major cellular pathways that were associated with very long-chain ceramides. These were mitochondria related pathways (e.g. ‘TCA cycle and respiratory electron transport’ and ‘Complex I biogenesis’), many terms involved in increased mRNA translation (e.g. Eukaryotic Translation Initiation’), overall cell cycle control and protein degradation (e.g. ‘SCF(Skp2)-mediated Degradation of p27/p21’) and stress related pathways (e.g. ‘Cellular response to chemical stress’ and ‘hypoxia’). (Fig. 4E and Supplemental table 4). While protein signatures associating with very long-chain ceramides reflected some of these trends, particularly increased translation, it also revealed notable differences. We saw a total absence of mitochondrial-related gene sets enriched with very long-chain ceramide abundance on a protein level (Fig. 4E). This was quite striking and suggested that increased mitochondrial gene expression is counterbalanced by elevated protein turnover, preventing an increase in mitochondrial protein abundance. This indicated some form of mitochondrial stress, where enhanced transcription did not translate to higher protein levels. Similarly, while transcriptomic data pointed to upregulation of stress-related pathways, including cell cycle regulation and cellular stress responses, these pathways were negatively associated at the protein level, suggesting a post-transcriptional suppression of their functional impact (Fig. 4E). These pathways were largely driven by proteasomal components, suggesting that while very long-chain ceramides were associated with a transcriptional response aimed at enhancing proteasomal function, this did not translate into increased proteasomal protein abundance. This discrepancy could indicate either a translational bottleneck or increased proteasomal turnover, ultimately limiting the accumulation of proteasomal subunits despite increased mRNA expression.

We noted that leptin gene expression was quantified in all muscle samples. Given that leptin is an adipocyte-specific gene that strongly correlates with overall adipocyte abundance (Zhang *et al*, 2002), we can use this as a marker of intermuscular adipocyte infiltration. We speculated that DAGs, but not other lipids quantified in our data, might correlate with *Lep* expression in muscle, since adipocytes typically have high concentrations of DAGs (Pradas *et al*, 2018). Not surprisingly, the total abundance of DAGs showed the strongest correlation with *Lep* expression (Fig. 4F), while very long-chain ceramides exhibited no such relationship (Fig. 4G). This further points towards the notion that DAGs were associated with IR as a product of obesity itself and probably as a consequence of contaminating adipocytes in the skeletal muscle.

## Discussion

Excessive lipid accumulation in skeletal muscle is a well-recognised cause of IR, with debate over whether DAGs or ceramides are the key intermediaries driving this pathology. We leveraged the natural genetic variation in DOz mice to examine how obesity shapes muscle lipid composition and systemic IR. This approach enabled comparisons across extreme IR states while controlling for obesity levels. We showed that both DAGs and ceramides were both linked to whole-body insulin sensitivity, but the association of DAGs was largely explained by the obesity state, while ceramides, particularly very long-chain ceramides, were elevated in the insulin resistant state independently of adiposity. These ceramide species were further linked with the activation of cellular stress response pathways and the regulation of mitochondrial proteostasis, both of which are closely linked to the pathophysiology of IR (Petersen *et al*, 2024; Greyslak *et al*, 2023; Diaz-Vegas *et al*, 2023). These findings suggest that while both DAGs and ceramides influence whole-body insulin sensitivity, ceramides may play a pivotal role in driving muscle IR.

Our observation that very long-chain ceramides (C20, C22, and C24) exhibited the greatest increase in the insulin-resistant group across any range of adiposity was surprising, as the long chain C18 ceramide have been implicated as the main ceramide in muscle linked to insulin resistance (Tonks *et al*, 2016; Bergman *et al*, 2016; Perreault *et al*, 2018). Despite this, some of the corroborating evidence for this claim is not straight forward. Knock out of CerS1, the enzyme thought to regulate C18 ceramide biosynthesis in skeletal muscle (Levy & Futerman, 2010), while leading to reduced muscle levels of C18 ceramide and protection from diet induced IR, appears to do so via an indirect mechanism involving FGF21 production (Turpin-Nolan *et al*, 2019). In fact, C18 ceramide is unlikely to drive muscle IR, as CerS1 inhibition depletes C18 ceramide while maintaining elevated very long-chain ceramide and this fails to restore insulin sensitivity in high-fat-fed mice (Turner *et al*, 2018). Regardless, other human studies have reported changes in very-long chain ceramides in insulin resistant people (Adams *et al*, 2004; Amati *et al*, 2011; Coen *et al*, 2010). Furthermore, LDL-bound C24:0 ceramides are elevated in plasma from people with diabetes and delivering lipoprotein-C24:0 ceramide conjugate to mice or myotubes induces muscle IR (Boon *et al*, 2013). C22 and C24 ceramides have also been implicated in inducing endoplasmic reticulum stress in muscle cells and tissue in a CerS2-dependent manner (McNally *et al*, 2022). Altogether, these data suggest that very long-chain ceramides may play a significant and underappreciated role in insulin resistance.

The above discussion poses the question as to what the relationship is between different ceramide species? Six ceramide synthases (CerS1-6) regulate *de novo* ceramide synthesis in mammals, producing a restrictive set of C14 to C36 ceramides (Turner *et al*, 2018). Long chain ceramides (C16 and C18) are often credited as having deleterious metabolic impact, whereas very long-chain ceramides, in particular C24, do not (Hammerschmidt & Brüning, 2022). These observations mainly stem from genetic mouse models, where either *Cers1*, *Cers2* or *Cers5/6* have been deleted. One of the confounding effects of many of these studies is that the reduction of one specific ceramide species is often accompanied by an elevation in the level of others. For example, *Cers1* null mice display depletion of C18 ceramide, but have elevated levels of C16- and C22-ceramides (Zhao *et al*, 2011; Ginkel *et al*, 2012; Turpin-Nolan *et al*, 2019). Similarly, decreasing CerS2 expression, which synthesises very chain-long ceramides, in mouse liver decreases C24:0 and increases C16 ceramides, which is thought to drive hepatic carcinoma (Bickert *et al*, 2018; Raichur *et al*, 2014). To complicate matters even more, ceramide levels with specific acyl chain lengths do not always align with CerS expression. For example, despite high CerS2 expression in the small intestine, C16- and C18-ceramides are the predominant species (Schiffmann *et al*, 2013). This suggests that factors beyond CerS expression and activity influence the acyl chain composition of ceramides in a given tissue. Ceramide production through alternative biosynthetic pathways, such as sphingomyelin hydrolysis and the salvage pathway, may markedly contribute to ceramide levels in skeletal muscle.

While elevated very long-chain ceramides were associated with a transcriptional response promoting cell cycle progression and stress adaptation, our proteomic data suggest increased protein turnover, possibly as a compensatory response to proteotoxic stress. Rather than simple translational repression, this pattern may reflect heightened protein degradation, ensuring the removal of damaged or misfolded proteins. In fact, ceramide accumulation has been linked with protein stability in both *in vitro* and *in vivo* studies (Diaz-Vegas *et al*, 2023; Greyslak *et al*, 2023; Turner *et al*, 2018). This could be explained by the increase in mitochondrial oxidative stress observed with ceramide overload. For example, ceramides can directly inhibit respiratory complex III leading to production of reaction oxygen species (Gudz *et al*, 1997), which in turn promotes mitochondrial proteostasis (Guarás *et al*, 2016). This suggests that elevated ceramides impose cellular stress that disrupts normal protein homeostasis, requiring active protein degradation to counteract potential proteotoxic effects.

Another question is how do ceramides drive IR? Mechanistically, ceramides can disrupt membrane fluidity and induce gel-to-fluid phase transitions and these properties are dependent on chain length and saturation status (Pinto *et al*, 2011). This is particularly important in mitochondria, where respiratory complexes are highly sensitive to membrane fluidity, and excessive gel-to-fluid transitions have been associated with respiratory deficiencies (Mileykovskaya & Dowhan, 2014). Furthermore, increased ceramides specifically in mitochondria are strongly linked with muscle IR in *in vitro* and *in vivo* studies (Petersen *et al*, 2024; Perreault *et al*, 2018; Diaz-Vegas *et al*, 2023; Greyslak *et al*, 2023). We propose that accumulation of very long-chain ceramides within mitochondria induces proteostatic and oxidative stress, driving muscle IR. This is supported by recent evidence demonstrating that mitochondrial ceramide levels, but not DAGs, are strong predictors of muscle IR regardless of the obesity state in humans (Petersen *et al*, 2024). However, determining how different lipid species impact mitochondrial membranes and respiratory complexes requires higher-resolution structural analyses (Mileykovskaya & Dowhan, 2014) and detailed subcellular lipid profiling (Perreault *et al*, 2018; Petersen *et al*, 2024).

Finally, we showed that leptin gene expression in skeletal muscle was highly associated with DAGs. The most likely explanation is that the primary contribution to elevated DAGs in muscle that are associated with adiposity, is the presence of infiltrating adipocytes in the muscle. In fact, magnetic resonance imaging has demonstrated a high level of adipocyte infiltration within skeletal muscle tissue in obesity and type 2 diabetes (Hilton *et al*, 2008). Similar observations have been made in mice fed a HFD (Khan *et al*, 2015). These adipocytes may arise from resident fibro-adipogenic progenitors, which have the capacity to differentiate into adipocytes and contribute to increased intermuscular lipid accumulation in obesity (Jia & Sowers, 2019). Regardless of their origin, adipocytes generally contain high concentrations of DAGs due to their primary role in lipid storage (Pradas *et al*, 2018). Therefore, even a smaller number of infiltrating adipocytes could significantly alter the muscle DAG profile.

In summary, our study links both DAGs and ceramides to whole-body IR, with DAGs driven by adiposity, and very long-chain ceramides potentially contributing as a cell-autonomous driver. While further research is needed to clarify the mechanisms behind this accumulation, our findings suggest that, beyond the current focus on C18 ceramides, targeting very long-chain ceramides could be a promising therapeutic strategy. Additionally, accounting for obesity will be crucial in future studies on muscle lipidome and whole-body metabolism.

## Author contributions: CRediT

Conceptualisation: SM, DEJ. Methodology: SM, JS, DEJ, GM. Formal Analysis, SM, CDR, HBC. Data curation: SM. Investigation: SM, HBC, KCC, MP, JK, SWCM, AH, JC, ADV, DEJ. Writing – Original Draft: SM, ADV. Writing – Review & Editing, All authors. Visualisation, SM. Supervision, DEJ; Funding Acquisition, DEJ.

## Acknowledgement

This work was supported by National Health and Medical Research Council (NHMRC) Project Grants: GNT1120201 and GNT1061122 to D.E.J.; GNT2013621 to D.E.J. and A.D.V; D.E.J. is an Australian Research Council (ARC) Laureate Fellow. The content is solely the responsibility of the authors and does not necessarily represent the official views of the NHMRC or ARC. The authors also acknowledge the facilities, and the scientific and technical assistance of Sydney Mass Spectrometry Facility, Laboratory Animal Services (both University of Sydney).

## Supplementary figure legends

**Figure S1.**
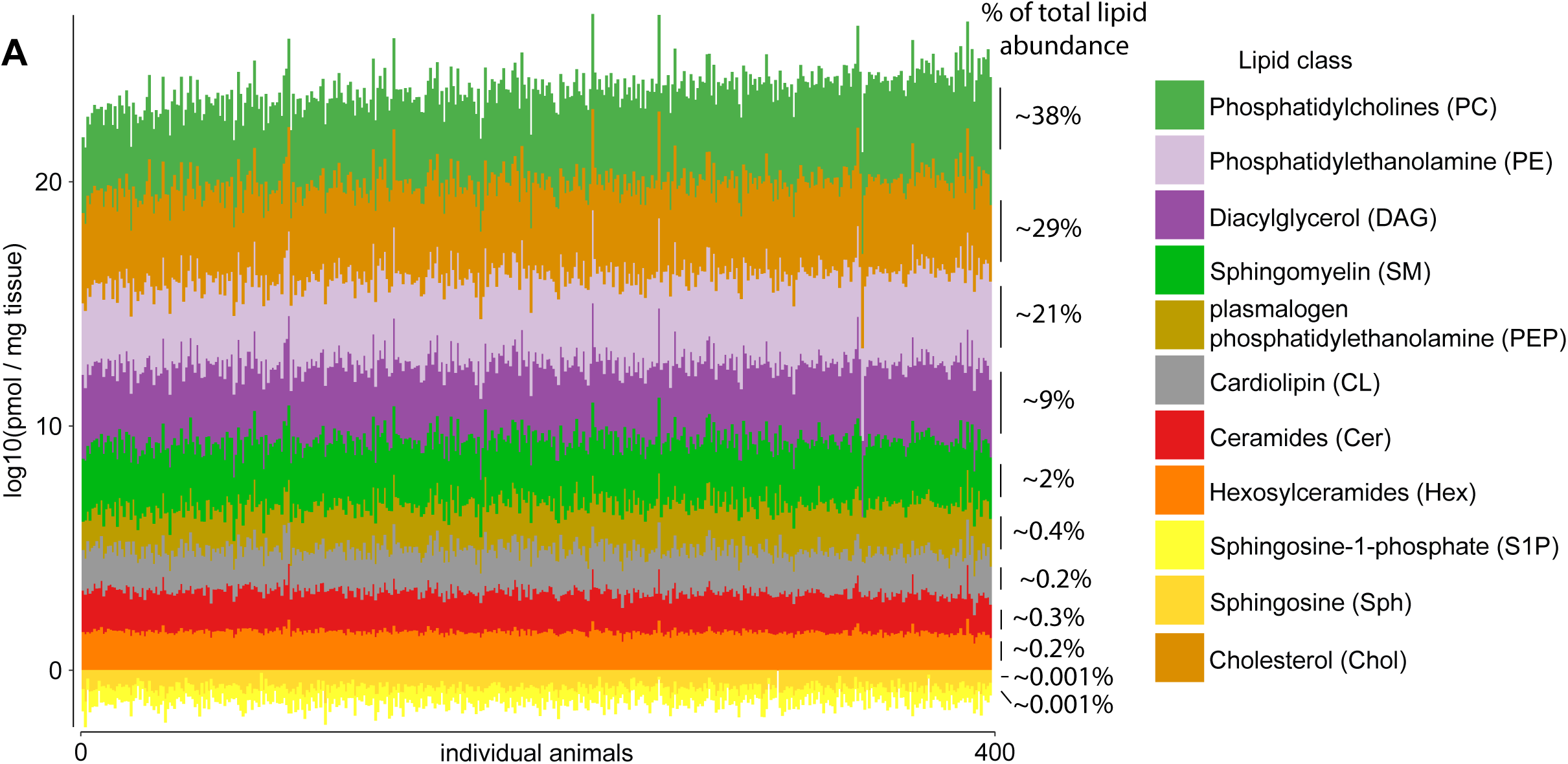
**A)** Absolute abundance of lipid classes quantified by targeted lipidomics ordered by abundance of absolute total phosphatidylcholine abundance. Estimates of total proportion by lipid class in percent are indicated to the right (relative mean of each class across 399 mice).

**Figure S2.**
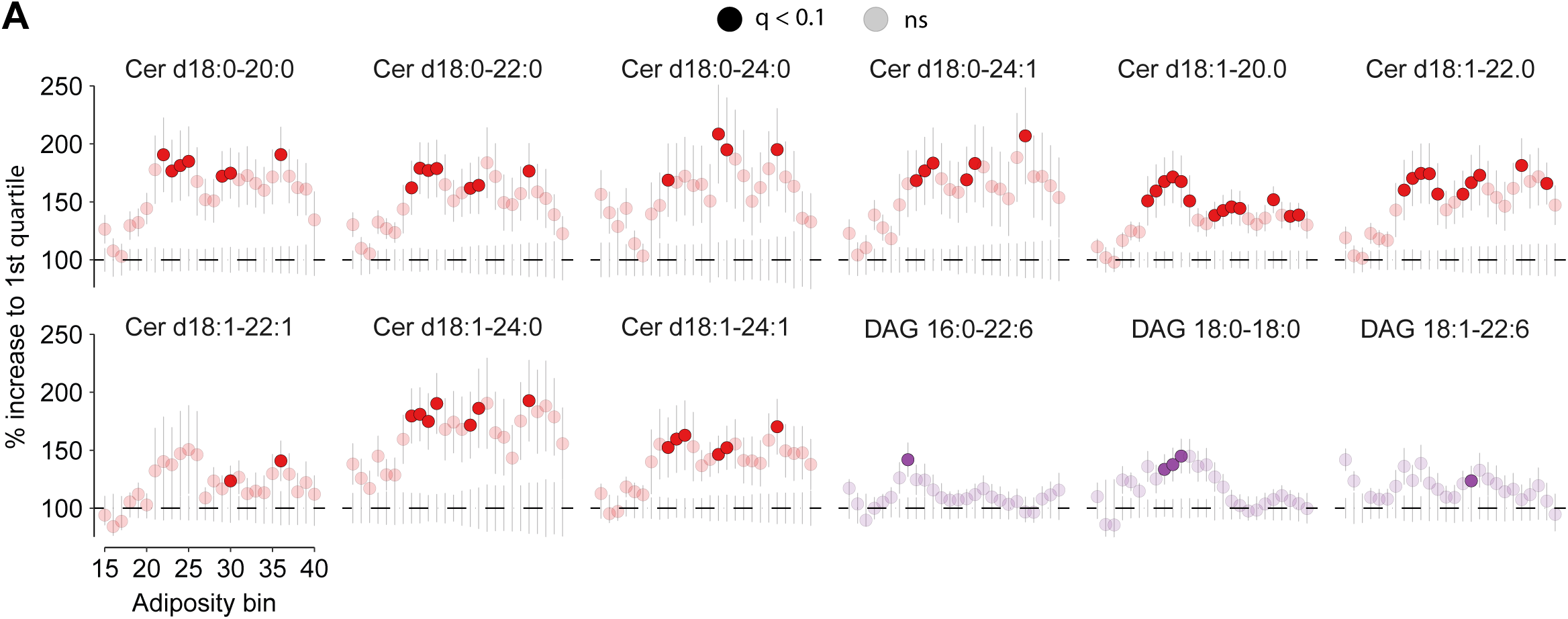
**A)** Comparison of ceramides (Cers) and diacylglycerols (DAGs) abundance between Q1 and Q4 relative to Q1 across all adiposity bins.

## REFERENCES

Adams JM 2nd, Pratipanawatr T, Berria R, Wang E, DeFronzo RA, Sullards MC & Mandarino LJ (2004) Ceramide content is increased in skeletal muscle from obese insulin-resistant humans. Diabetes 53: 25–31

Amati F, Dubé JJ, Alvarez-Carnero E, Edreira MM, Chomentowski P, Coen PM, Switzer GE, Bickel PE, Stefanovic-Racic M, Toledo FGS, et al (2011) Skeletal muscle triglycerides, diacylglycerols, and ceramides in insulin resistance: another paradox in endurance-trained athletes? Diabetes 60: 2588–2597

Beckerman M, Harel C, Michael I, Klip A, Bilan PJ, Gallagher EJ, LeRoith D, Lewis EC, Karnieli E & Levenberg S (2021) GLUT4-overexpressing engineered muscle constructs as a therapeutic platform to normalize glycemia in diabetic mice. Sci Adv 7: eabg3947

Bergman BC, Brozinick JT, Strauss A, Bacon S, Kerege A, Bui HH, Sanders P, Siddall P, Wei T, Thomas MK, et al (2016) Muscle sphingolipids during rest and exercise: a C18:0 signature for insulin resistance in humans. Diabetologia 59: 785–798

Bickert A, Kern P, van Uelft M, Herresthal S, Ulas T, Gutbrod K, Breiden B, Degen J, Sandhoff K, Schultze JL, et al (2018) Inactivation of ceramide synthase 2 catalytic activity in mice affects transcription of genes involved in lipid metabolism and cell division. Biochim Biophys Acta Mol Cell Biol Lipids 1863: 734–749

Boon J, Hoy AJ, Stark R, Brown RD, Meex RC, Henstridge DC, Schenk S, Meikle PJ, Horowitz JF, Kingwell BA, et al (2013) Ceramides contained in LDL are elevated in type 2 diabetes and promote inflammation and skeletal muscle insulin resistance. Diabetes 62: 401–410

Bray NL, Pimentel H, Melsted P & Pachter L (2016) Near-optimal probabilistic RNA-seq quantification. Nat Biotechnol 34: 525–527

Chaurasia B & Summers SA (2021) Ceramides in Metabolism: Key Lipotoxic Players. Annu Rev Physiol 83: 303–330

Chella Krishnan K, El Hachem E-J, Keller MP, Patel SG, Carroll L, Vegas AD, Gerdes Gyuricza I, Light C, Cao Y, Pan C, et al (2023) Genetic architecture of heart mitochondrial proteome influencing cardiac hypertrophy. Elife 12

Churchill GA, Gatti DM, Munger SC & Svenson KL (2012) The Diversity Outbred mouse population. Mamm Genome 23: 713–718

Coen PM, Dubé JJ, Amati F, Stefanovic-Racic M, Ferrell RE, Toledo FGS & Goodpaster BH (2010) Insulin resistance is associated with higher intramyocellular triglycerides in type I but not type II myocytes concomitant with higher ceramide content. Diabetes 59: 80–88

Cohen DE & Fisher EA (2013) Lipoprotein metabolism, dyslipidemia, and nonalcoholic fatty liver disease. Semin Liver Dis 33: 380–388

Després JP, Lamarche B, Mauriège P, Cantin B, Dagenais GR, Moorjani S & Lupien PJ (1996) Hyperinsulinemia as an independent risk factor for ischemic heart disease. N Engl J Med 334: 952–957

Diaz-Vegas A, Cooke KC, Cutler HB, Yau B, Masson SWC, Harney D, Fuller OK, Potter M, Madsen S, Craw NR, et al (2024a) Deletion of miPEP in adipocytes protects against obesity and insulin resistance by boosting muscle metabolism. Mol Metab: 101983

Diaz-Vegas A, Madsen S, Cooke KC, Carroll L, Khor JXY, Turner N, Lim XY, Astore MA, Morris JC, Don AS, et al (2023) Mitochondrial electron transport chain, ceramide, and coenzyme Q are linked in a pathway that drives insulin resistance in skeletal muscle. Elife 12

Diaz-Vegas AR, Don AS & Burchfield JG (2024b) Analysis and quantification of the mitochondrial-ER lipidome. Bio Protoc 14: e5028

Erion DM & Shulman GI (2010) Diacylglycerol-mediated insulin resistance. Nat Med 16: 400–402

Ewels P, Magnusson M, Lundin S & Käller M (2016) MultiQC: summarize analysis results for multiple tools and samples in a single report. Bioinformatics 32: 3047– 3048

van Gerwen J, Masson SWC, Cutler HB, Vegas AD, Potter M, Stöckli J, Madsen S, Nelson ME, Humphrey SJ & James DE (2024) The genetic and dietary landscape of the muscle insulin signalling network. Elife 12

Ginkel C, Hartmann D, vom Dorp K, Zlomuzica A, Farwanah H, Eckhardt M, Sandhoff R, Degen J, Rabionet M, Dere E, et al (2012) Ablation of neuronal ceramide synthase 1 in mice decreases ganglioside levels and expression of myelin-associated glycoprotein in oligodendrocytes. J Biol Chem 287: 41888–41902

Green CD, Maceyka M, Cowart LA & Spiegel S (2021) Sphingolipids in metabolic disease: The good, the bad, and the unknown. Cell Metab 33: 1293–1306

Greyslak KT, Hetrick B, Bergman BC, Dean TA, Wesolowski SR, Gannon M, Schenk S, Sullivan EL, Aagaard KM, Kievit P, et al (2023) A Maternal Western-Style Diet Impairs Skeletal Muscle Lipid Metabolism in Adolescent Japanese Macaques. Diabetes 72: 1766–1780

Guarás A, Perales-Clemente E, Calvo E, Acín-Pérez R, Loureiro-Lopez M, Pujol C, Martínez-Carrascoso I, Nuñez E, García-Marqués F, Rodríguez-Hernández MA, et al (2016) The CoQH2/CoQ Ratio Serves as a Sensor of Respiratory Chain Efficiency. Cell Rep 15: 197–209

Gudz TI, Tserng KY & Hoppel CL (1997) Direct inhibition of mitochondrial respiratory chain complex III by cell-permeable ceramide. J Biol Chem 272: 24154–24158

Hammerschmidt P & Brüning JC (2022) Contribution of specific ceramides to obesity-associated metabolic diseases. Cell Mol Life Sci 79: 395

Hilton TN, Tuttle LJ, Bohnert KL, Mueller MJ & Sinacore DR (2008) Excessive adipose tissue infiltration in skeletal muscle in individuals with obesity, diabetes mellitus, and peripheral neuropathy: association with performance and function. Phys Ther 88: 1336–1344

Hunter DJ (2005) Gene-environment interactions in human diseases. Nat Rev Genet 6: 287–298

James DE, Stöckli J & Birnbaum MJ (2021) The aetiology and molecular landscape of insulin resistance. Nat Rev Mol Cell Biol 22: 751–771

Jani S, Da Eira D, Hadday I, Bikopoulos G, Mohasses A, de Pinho RA & Ceddia RB (2021) Distinct mechanisms involving diacylglycerol, ceramides, and inflammation underlie insulin resistance in oxidative and glycolytic muscles from high fat-fed rats. Sci Rep 11: 19160

Jia G & Sowers JR (2019) Increased fibro-adipogenic progenitors and intramyocellular lipid accumulation in obesity-related skeletal muscle dysfunction. Diabetes 68: 18– 20

Khan IM, Perrard XY, Brunner G, Lui H, Sparks LM, Smith SR, Wang X, Shi Z-Z, Lewis DE, Wu H, et al (2015) Intermuscular and perimuscular fat expansion in obesity correlates with skeletal muscle T cell and macrophage infiltration and insulin resistance. Int J Obes (Lond) 39: 1607–1618

Kim JK, Fillmore JJ, Chen Y, Yu C, Moore IK, Pypaert M, Lutz EP, Kako Y, Velez-Carrasco W, Goldberg IJ, et al (2001) Tissue-specific overexpression of lipoprotein lipase causes tissue-specific insulin resistance. Proc Natl Acad Sci U S A 98: 7522–7527

Koves TR, Ussher JR, Noland RC, Slentz D, Mosedale M, Ilkayeva O, Bain J, Stevens R, Dyck JRB, Newgard CB, et al (2008) Mitochondrial overload and incomplete fatty acid oxidation contribute to skeletal muscle insulin resistance. Cell Metab 7: 45–56

Kraegen EW, Bruce C, Hegarty BD, Ye J-M, Turner N & Cooney G (2009) AMP-activated protein kinase and muscle insulin resistance. Front Biosci (Landmark Ed) 14: 4658–4672

Langfelder P & Horvath S (2008) WGCNA: an R package for weighted correlation network analysis. BMC Bioinformatics 9: 559

Larsen PJ & Tennagels N (2014) On ceramides, other sphingolipids and impaired glucose homeostasis. Mol Metab 3: 252–260

Levy M & Futerman AH (2010) Mammalian ceramide synthases. IUBMB Life 62: 347– 356

Masson SWC, Cutler HB & James DE (2024) Unlocking metabolic insights with mouse genetic diversity. EMBO J 43: 4814–4821

Masson SWC, Madsen S, Cooke KC, Potter M, Vegas AD, Carroll L, Thillainadesan S, Cutler HB, Walder KR, Cooney GJ, et al (2023) Leveraging genetic diversity to identify small molecules that reverse mouse skeletal muscle insulin resistance. Elife 12

McNally BD, Ashley DF, Hänschke L, Daou HN, Watt NT, Murfitt SA, MacCannell ADV, Whitehead A, Bowen TS, Sanders FWB, et al (2022) Long-chain ceramides are cell non-autonomous signals linking lipotoxicity to endoplasmic reticulum stress in skeletal muscle. Nat Commun 13: 1748

Mileykovskaya E & Dowhan W (2014) Cardiolipin-dependent formation of mitochondrial respiratory supercomplexes. Chem Phys Lipids 179: 42–48

Morino K, Petersen KF & Shulman GI (2006) Molecular mechanisms of insulin resistance in humans and their potential links with mitochondrial dysfunction. Diabetes 55 Suppl 2: S9–S15

Nelson ME, Madsen S, Cooke KC, Fritzen AM, Thorius IH, Masson SWC, Carroll L, Weiss FC, Seldin MM, Potter M, et al (2022) Systems-level analysis of insulin action in mouse strains provides insight into tissue-and pathway-specific interactions that drive insulin resistance. Cell Metab 34: 227–239.e6

Perreault L, Newsom SA, Strauss A, Kerege A, Kahn DE, Harrison KA, Snell-Bergeon JK, Nemkov T, D’Alessandro A, Jackman MR, et al (2018) Intracellular localization of diacylglycerols and sphingolipids influences insulin sensitivity and mitochondrial function in human skeletal muscle. JCI Insight 3

Petersen MC, Smith GI, Palacios HH, Farabi SS, Yoshino M, Yoshino J, Cho K, Davila-Roman VG, Shankaran M, Barve RA, et al (2024) Cardiometabolic characteristics of people with metabolically healthy and unhealthy obesity. Cell Metab 36: 745– 761.e5

Pinto SN, Silva LC, Futerman AH & Prieto M (2011) Effect of ceramide structure on membrane biophysical properties: the role of acyl chain length and unsaturation. Biochim Biophys Acta 1808: 2753–2760

Pradas I, Huynh K, Cabré R, Ayala V, Meikle PJ, Jové M & Pamplona R (2018) Lipidomics reveals a tissue-specific fingerprint. Front Physiol 9: 1165

Raichur S, Wang ST, Chan PW, Li Y, Ching J, Chaurasia B, Dogra S, Öhman MK, Takeda K, Sugii S, et al (2014) CerS2 haploinsufficiency inhibits β-oxidation and confers susceptibility to diet-induced steatohepatitis and insulin resistance. Cell Metab 20: 687–695

Robinson MD, McCarthy DJ & Smyth GK (2010) edgeR: a Bioconductor package for differential expression analysis of digital gene expression data. Bioinformatics 26: 139–140

Schiffmann S, Birod K, Männich J, Eberle M, Wegner M-S, Wanger R, Hartmann D, Ferreiros N, Geisslinger G & Grösch S (2013) Ceramide metabolism in mouse tissue. Int J Biochem Cell Biol 45: 1886–1894

Soneson C, Love MI & Robinson MD (2015) Differential analyses for RNA-seq: transcript-level estimates improve gene-level inferences. F1000Res 4: 1521

Steneberg P, Sykaras AG, Backlund F, Straseviciene J, Söderström I & Edlund H (2015) Hyperinsulinemia enhances hepatic expression of the fatty acid transporter Cd36 and provokes hepatosteatosis and hepatic insulin resistance. J Biol Chem 290: 19034–19043

Strimmer K (2008) fdrtool: a versatile R package for estimating local and tail area-based false discovery rates. Bioinformatics 24: 1461–1462

Thiebaud D, Jacot E, DeFronzo RA, Maeder E, Jequier E & Felber JP (1982) The effect of graded doses of insulin on total glucose uptake, glucose oxidation, and glucose storage in man. Diabetes 31: 957–963

Tonks KT, Coster AC, Christopher MJ, Chaudhuri R, Xu A, Gagnon-Bartsch J, Chisholm DJ, James DE, Meikle PJ, Greenfield JR, et al (2016) Skeletal muscle and plasma lipidomic signatures of insulin resistance and overweight/obesity in humans. Obesity (Silver Spring) 24: 908–916

Tsao TS, Burcelin R, Katz EB, Huang L & Charron MJ (1996) Enhanced insulin action due to targeted GLUT4 overexpression exclusively in muscle. Diabetes 45: 28–36

Turner N, Lim XY, Toop HD, Osborne B, Brandon AE, Taylor EN, Fiveash CE, Govindaraju H, Teo JD, McEwen HP, et al (2018) A selective inhibitor of ceramide synthase 1 reveals a novel role in fat metabolism. Nat Commun 9: 3165

Turpin-Nolan SM, Hammerschmidt P, Chen W, Jais A, Timper K, Awazawa M, Brodesser S & Brüning JC (2019) CerS1-Derived C18:0 Ceramide in Skeletal Muscle Promotes Obesity-Induced Insulin Resistance. Cell Rep 26: 1–10.e7

Wang L, Valencak TG & Shan T (2024) Fat infiltration in skeletal muscle: Influential triggers and regulatory mechanism. iScience 27: 109221

Wu T, Hu E, Xu S, Chen M, Guo P, Dai Z, Feng T, Zhou L, Tang W, Zhan L, et al (2021) clusterProfiler 4.0: A universal enrichment tool for interpreting omics data. Innovation (Camb) 2: 100141

Yu G & He Q-Y (2016) ReactomePA: an R/Bioconductor package for reactome pathway analysis and visualization. Mol Biosyst 12: 477–479

Zhang AMY, Xia YH, Lin JSH, Chu KH, Wang WCK, Ruiter TJJ, Yang JCC, Chen N, Chhuor J, Patil S, et al (2023) Hyperinsulinemia acts via acinar insulin receptors to initiate pancreatic cancer by increasing digestive enzyme production and inflammation. Cell Metab 35: 2119–2135.e5

Zhang Y, Guo K-Y, Diaz PA, Heo M & Leibel RL (2002) Determinants of leptin gene expression in fat depots of lean mice. Am J Physiol Regul Integr Comp Physiol 282: R226–34

Zhao L, Spassieva SD, Jucius TJ, Shultz LD, Shick HE, Macklin WB, Hannun YA, Obeid LM & Ackerman SL (2011) A deficiency of ceramide biosynthesis causes cerebellar purkinje cell neurodegeneration and lipofuscin accumulation. PLoS Genet 7: e1002063

Zierath JR (2007) The path to insulin resistance: paved with ceramides? Cell Metab 5: 161–163

